# Risk factors associated with loss of hepatitis B virus surface antibody in patients with HBV surface-antigen negative/surface antibody positive serostatus receiving biologic DMARDs to treat rheumatic diseases – a nested case-control study

**DOI:** 10.1101/661850

**Authors:** Ming-Hui Hung, Ya-Chih Tien, Ying-Ming Chiu

## Abstract

**Objectives:** To elucidate risk factors for loss of hepatitis B virus (HBV) surface antibody (anti-HBs) in patients with rheumatic diseases and HBV surface-antigen negative/anti-HBs positive (HBsAg−/anti-HBs+) serostatus during biologic disease-modifying anti-rheumatic drug (DMARD) treatment.

**Methods:** This nested case-control study prospectively enrolled patients with rheumatoid arthritis, ankylosing spondylitis, psoriatic arthritis/psoriasis, and juvenile idiopathic arthritis, who were treated with biologic DMARDs from January 2013 to September 2017. The analytic sample included patients with HBsAg−/anti-HBs+ serostatus. Anti-HBs titers were monitored, and cases defined as anti-HBs <10 mIU/mL during follow-up. Cases were matched one-to-all with controls with anti-HBs ≥10 mIU/mL on the same event date and equivalent durations of biologic DMARDs treatment. Between-group characteristics were compared and risk factors for anti-HBs loss elucidated by conditional logistic regression analyses.

**Results:** Among 189 enrolled patients, 15 cases were matched with 211 controls. Risk factors associated with anti-HBs loss in multivariate analysis were low baseline anti-HBs titer (adjusted risk ratio = 0.96, 95% CI 0.93–0.99) and chronic kidney disease (adjusted risk ratio = 26.25, 95% CI 1.85–372.35). All cases had baseline anti-HBs titer <100 mIU/mL, and none developed HBV reactivation upon losing anti-HBs.

**Conclusions:** In addition to low baseline anti-HBs titer, chronic kidney disease is also an independent risk factors associated with loss of anti-HBs in patients with HBsAg−/anti-HBs+ serostatus who receive biologic DMARDs to treat rheumatic diseases.

**Significance:** Given that loss of anti-HBs precedes HBV reactivation and that the use of biologic DMARDs is increasingly widespread nowadays, understanding those who are at risk of loss of anti-HBs is an important and practical clinical issue.

**Innovation:** In addition to low baseline anti-HBs titer, chronic kidney disease is also an independent risk factors associated with loss of anti-HBs in patients with HBsAg−/anti-HBs+ serostatus who receive biologic DMARDs to treat rheumatic diseases.

## INTRODUCTION

Hepatitis B virus (HBV) infection is a major public health concern worldwide. Hepatitis B reactivation is characterized by HBV replication and the recurrence of active necro-inflammatory liver disease. HBV reactivation after chemotherapy or immunosuppressive therapy, both in people with HBV surface-antigen positive (HBsAg+) serostatus and those who are HBsAg-negative with antibodies against HBV core-antigen or surface-antigen (HBsAg−/anti-HBc+ or anti-HBs+),[1–3], is an increasingly recognized problem,[1, 4] because reactivation can interrupt the treatment of underlying disease,[5] and may presage severe hepatitis or death.

Manifestation of serum HBV DNA (viremia) is widely acknowledged to be an important definition of HBV reactivation.[6] However, clinical HBV reactivation is not an inevitable consequence of HBV DNA viremia, which can be transient, especially whilst anti-HBs status is still positive.[7] Furthermore, in cases of manifest HBV viremia, anti-HBs loss is a major determinant of, and almost precedes, HBV reactivation.[7, 8]

Because anti-HBs loss is known to occur after immunosuppressive therapy [9, 10] and almost always precedes HBV reactivation,[11] risk factors associated with anti-HBs negativity (<10 mIU/ml) are particularly important, especially given burgeoning use of tumor necrosis factor inhibitors (anti-TNF) and other biologic disease-modifying anti-rheumatic drugs (DMARDs) to treat various autoimmune diseases, and growing evidence of elevated HBV reactivation rates in this setting.[1, 12]

However, the risk factors of anti-HBs loss in rheumatic patients undergoing biologic DMARDs therapy is unknown. Hence, we conducted a nested case-control study in a prospective cohort of hospital patients to investigate this research question.

## METHODS

### Study subjects

The study population comprised patients at Changhua Christian Hospital, Taiwan, with rheumatoid arthritis, ankylosing spondylitis, psoriasis, psoriatic arthritis, and juvenile idiopathic arthritis, who were treated with biologic DMARDs from January 2013 to September 2017. Only patients with HBsAg−/anti-HBs+ serostatus were enrolled (Figure 1); subjects with HBsAg+ or HBsAg−/anti-HBs− serostatus were excluded. All enrolled participants fulfilled international diagnostic criteria for these diseases and were treated in accordance with national consensus recommendations for screening and management of viral hepatitis,[13] which recommend HBV serology tests and HBV DNA monitoring every 6 months.

**Figure 1.**
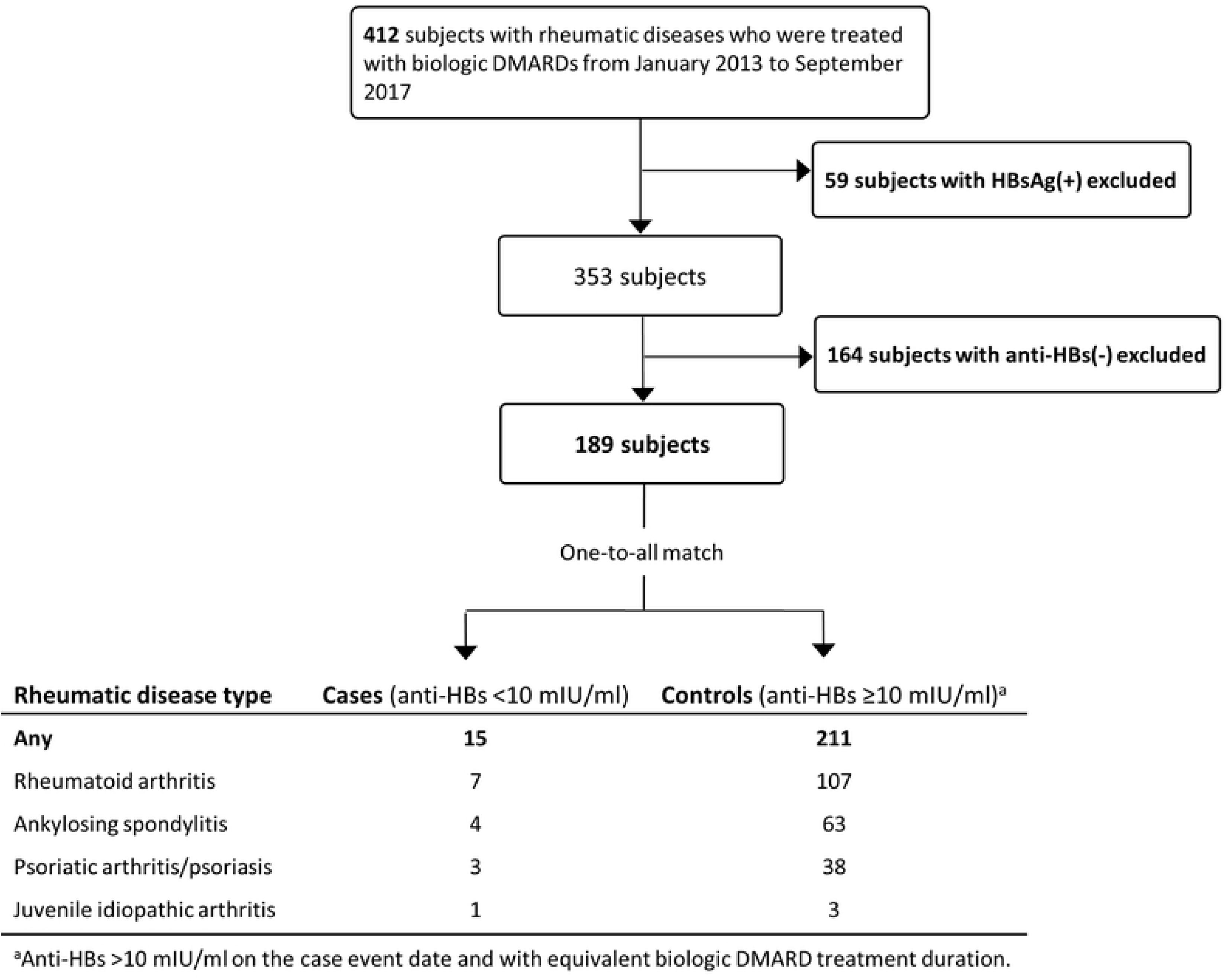
Case-control selection flow chart. DMARDs, disease-modifying anti-rheumatic drugs; HBV, hepatitis B virus, DNA, deoxyribonucleic acid; anti-HBs, HBV surface antibody; HBsAg, HBV surface-antigen; mIU, million International Units.

### Hepatitis B serologic testing and HBV DNA

HBV assays included serum HBsAg, anti-HBs and anti-HBc, measured by Architect i2000SR chemiluminescent microparticle immunoassay (Abbott Laboratories, Abbot Park, Illinois, USA). Serum HBV DNA viral load was quantified by Abbott RealTime HBV (Abbott Laboratories, Abbott Park, Illinois, USA), with a minimal sensitivity of 10 IU/ml.

### Covariate information

Baseline data included: age, sex, type of rheumatic disease (rheumatoid arthritis, ankylosing spondylitis, psoriatic arthritis/psoriasis, juvenile idiopathic arthritis), accumulated doses of conventional DMARDs (prednisolone, hydroxychloroquine, sulfasalazine, methotrexate, leflunomide, cyclosporine) and biologic DMARDs (etanercept, adalumumab, golimumab, ustekimumab, tocitizumab, rituximab, abatacept, tofacitinib). Chronic kidney disease was defined as estimated glomerular filtration rate <60 mL/min/1.73 m2. Chronic liver disease status, including fatty liver and parenchymal liver disease, was determined from medical charts or hepatic ultrasound results.

### Nested case-control design

Due to the complexity and varying durations of drug exposures in this study population, we used a nested case-control design, which is a valid alternative to cohort analysis that does not compromise statistical power.[14, 15] Cases were defined upon occurrences of serum anti-HBs titer <10 mIU/mL during follow-up, and the date that anti-HBs loss was ascertained designated the event date. Each case was matched one-to-all with subjects whose serum HBsAb was ≥10 mIU/ml on the respective case event date and who had an equivalent duration of biologic DMARDs treatment. One patient could therefore serve as a control repeatedly during follow-up, albeit at different times, and control subjects could become cases during the study.[16]

### Statistics

All analyses were performed using SAS® software, Version 9.2 for Windows (SAS Institute Inc., Cary, NC, USA); p-value <0.05 for two-sided tests was considered statistically significant. Continuous variables were expressed as means plus/minus standard deviation or median [range], categorical variables as numbers (percentages). Conditional logistic regression analysis was used to estimate risk ratios and 95% confidence intervals for loss of anti-HBs; putative associated factors included age, sex, type of rheumatic disease, traditional DMARDs, biologic DMARDs (anti-TNF or others), comorbidity, and baseline anti-HBs.

## RESULTS

### Demographic characteristics and clinical status

The analytic sample comprised 15 cases and 211 matched controls (Figure 1); Table 1 shows their demographic and clinical characteristics.

**Table 1.** Baseline characteristics.

| Data show mean ± standard deviation, median [range], or number (%) | Cases | Controls |
| --- | --- | --- |
| Number | 15 | 221 |
| Age (years) | 46.0 ± 18.9 | 47.2 ± 13.1 |
| Sex |  |  |
| Female | 8 (53.3%) | 100 (47.4%) |
| Male | 7 (46.7%) | 111 (52.6%) |
| Rheumatic disease |  |  |
| Rheumatoid arthritis | 7 (46.7%) | 107 (50.7%) |
| Ankylosing spondylitis | 4 (26.7%) | 63 (29.9%) |
| Psoriatic arthritis/Psoriasis | 3 (20%) | 38 (18.0%) |
| Juvenile idiopathic arthritis | 1 (6.7%) | 3 (1.4%) |
| Baseline serum anti-HBs titer (mIU/ml) | 17.3 [10.1–64.7] | 136.7 [11.5–1010]<sup>a</sup> |
| Biologic DMARDs |  |  |
| Anti-TNF (Etanercept, Adalimumab, Golimumab) | 10 (66.7%) | 173 (82.0%) |
| Not anti-TNF | 5 (33.3%) | 38 (18.0%) |
| Abatacept | 0 | 7 (3.3%) |
| Rituximab | 1 (6.7%) | 3 (1.4%) |
| Tocilizumab | 2 (13.3%) | 7 (3.3%) |
| Tofacitinib | 0 | 2 (0.9%) |
| Ustekinumab | 2 (13.3%) | 19 (9.0%) |
| Conventional DMARDs (accumulated dose) |  |  |
| Methotrexate (mg) | 297 ± 444 | 452 ± 628 |
| Leflunomide (mg) | 1052 ± 2532 | 276 ± 1593 |
| Sulfasalazine (g) | 393 ± 393 | 185 ± 272 |
| Hydroxychloroquine (g) | 44 ± 87 | 40 ± 78 |
| Cyclosporine (g) | 14 ± 26 | 8 ± 25 |
| Steroid (accumulated dose, mg) | 1807 ± 2635 | 1282 ± 1789 |
| Comorbidities |  |  |
| Hepatitis C virus antibody positive | 2 (13.3%) | 2 (1.0%) |
| Diabetes mellitus | 4 (26.7%) | 7 (3.3%) |
| Chronic liver disease | 5 (33.3%) | 27 (12.8%) |
| Chronic kidney disease | 2 (13.3%) | 2 (1.0%) |

Anti-HBs, hepatitis B virus surface antibody; DMARD, disease-modifying anti-rheumatic drug; TNF, tumor necrosis factor.
<sup>a</sup>Maximal detectable limit.

Mean age, sex ratio, and rheumatic disease types were similar between case and control groups. Compared with controls, cases had lower baseline serum anti-HBs titer, more prevalent comorbidities (including hepatitis C infection, chronic liver disease, diabetes mellitus, chronic kidney disease), and relatively higher accumulated doses of sulfasalazine, leflunomide, and steroids, but a lower accumulated dose of methotrexate. Most subjects in both groups used anti-TNF agents (etanercept, adalimumab, golimumab).

### Risk factors for anti-HBs loss

Table 2 shows risk factors associated with loss of anti-HBs in conditional logistic regression analyses. The only factors remaining significant in the multivariate model, were low baseline serum anti-HBs titer and chronic kidney disease.

**Table 2.** Risk factors associated with loss of anti-HBs.

|  | Univariate analysis |  | Multivariate analysis |  |
| --- | --- | --- | --- | --- |
|  | Risk ratio (95% CI) | *p*-value | Risk ratio (95% CI) | *p*-value |
| Age | 0.99 (0.96–1.03) | 0.72 |  |  |
| Sex |  |  |  |  |
| Female | 1 (reference) | 0.60 |  |  |
| Male | 0.76 (0.27–2.15) |  |  |  |
| Rheumatic disease |  |  |  |  |
| Rheumatoid arthritis | 1 (reference) | 0.59 |  |  |
| Ankylosing spondylitis | 0.93 (0.27–3.24) |  |  |  |
| Psoriatic arthritis/Psoriasis | 1.13 (0.28–4.47) |  |  |  |
| Juvenile idiopathic arthritis | 4.83 (0.47–49.98) |  |  |  |
| Baseline serum anti-HBs | 0.96 (0.94–0.99) | 0.01 | 0.96 (0.93–0.99) | 0.01 |
| Biologic DMARDs |  |  |  |  |
| Anti-TNF | 1 (reference) | 0.21 |  |  |
| Not anti-TNF | 2.12 (0.66–6.83) |  |  |  |
| Conventional DMARDs |  |  |  |  |
| Methotrexate | 0.54 (0.18–1.61) | 0.27 |  |  |
| Sulfasalazine | 1.20 (1.03–1.38) | 0.02 |  |  |
| Hydroxychloroquine | 1.07 (0.55–2.08) | 0.84 |  |  |
| Cyclosporine | 1.07 (0.91–1.26) | 0.40 |  |  |
| Leflunomide | 1.18 (0.97–1.43) | 0.11 |  |  |
| Steroid | 1.15 (0.89–1.50) | 0.29 |  |  |
| Comorbidity |  |  |  |  |
| Hepatitis C virus antibody positive | 19.62 (1.70–226.88) | 0.02 |  |  |
| Diabetes mellitus | 9.26 (2.41–35.51) | 0.01 |  |  |
| Chronic liver disease | 3.32 (1.05–10.48) | 0.04 |  |  |
| Chronic kidney disease | 12.33 (1.71–88.93) | 0.01 | 26.25 (1.85–372.35) | 0.02 |

Anti-HBs, hepatitis B virus surface antibody; DMARD, disease-modifying anti-rheumatic drug; TNF, tumor necrosis factor.

### Clinical features and outcomes of subjects with anti-HBs loss

Seven of the 15 cases had rheumatoid arthritis (Table 3). All cases’ baseline anti-HBs titers were <100 mIU/mL. Ten cases were prescribed anti-TNF agents: four etanercept, four adalimumab, two golimumab. Two cases each were prescribed ustekinumab or tocilizumab. Only one case received rituximab. Serum HBV DNA upon anti-HBs loss was checked in 11/15 cases and only one had a detectable viral load. No cases developed HBV reactivation, had alanine transaminase elevation, or received any anti-viral treatment.

**Table 3.**
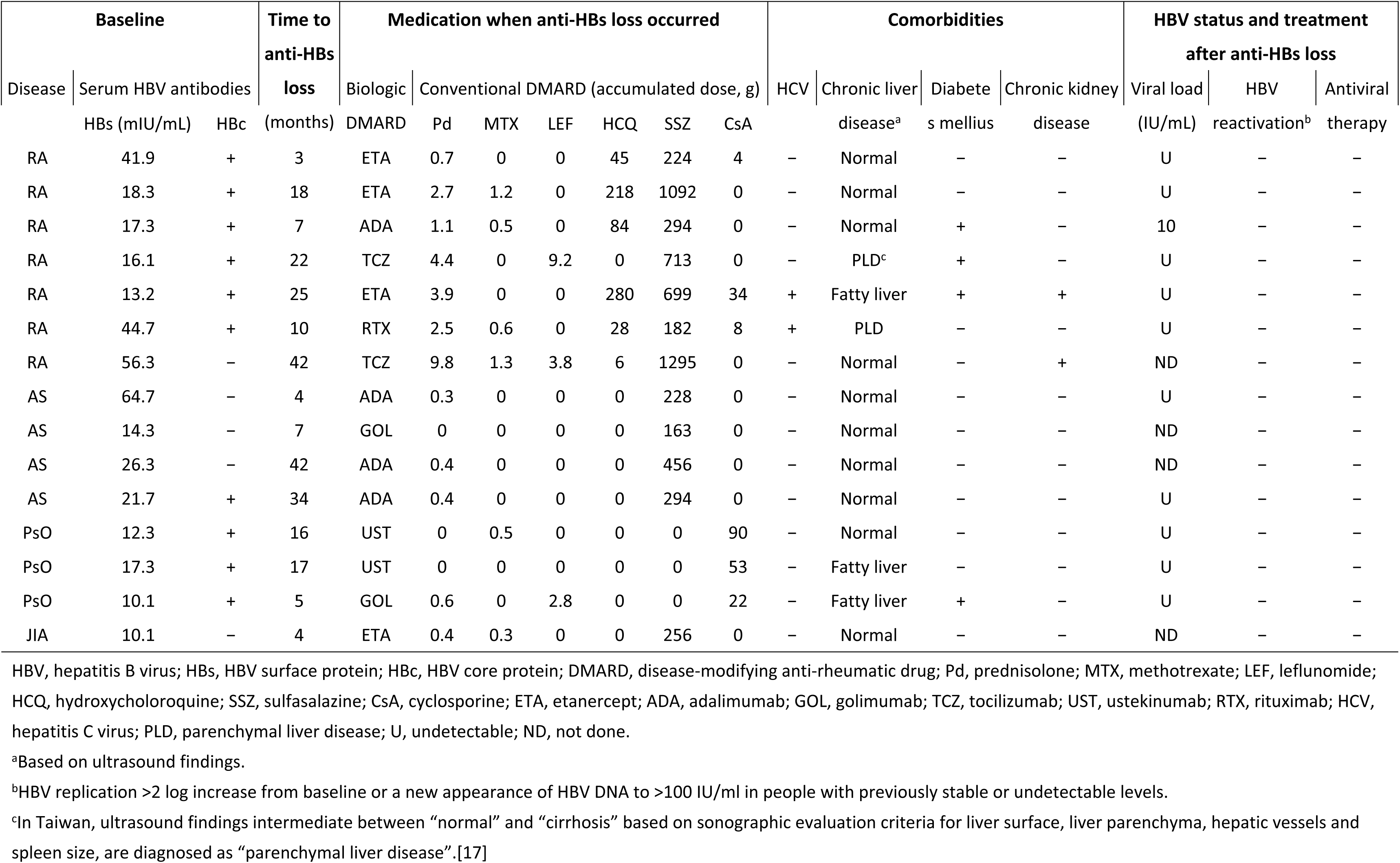
Characteristics of biologic DMARDs-treated patients with anti-HBs loss.

## DISCUSSION

To best of our knowledge, this is the first reported investigation of risk factors associated with loss of anti-HBs in rheumatic patients during biologic DMARDs therapy, after controlling for putative risk factors. We discovered that low baseline anti-HBs level and chronic kidney disease were significantly associated with anti-HBs loss.

In cases of manifest HBV viremia, anti-HBs loss is a major determinant of, and almost precedes, HBV reactivation. Although anti-HBs is important in protecting against HBV reactivation,[1, 8] our results demonstrate that this protective power is easily lost in cases of low baseline anti-HBs titer. Previous guidelines or reviews have propounded anti-HBs testing in baseline screening prior to using biologic DMARDs, because patients with baseline anti-HBs+ serostatus have lower risk of HBV reactivation.[6, 12, 18] However, current guidelines, particularly those focused on biologic DMARDs users, neither describe nor elucidate the potential risk of anti-HBs loss during biologic DMARDs therapy.[1, 12, 19] We found baseline anti-HBs titer <100 mU/mL to increase the risk of subsequent anti-HBs loss during biologic DMARDs therapy, despite anti-HBs+ status at baseline. Likewise, lymphoma patients receiving rituximab-based chemotherapy had a similar cut-off titer for risk of anti-HBs loss.[20] These results imply that clinicians should closely monitor patients with low baseline anti-HBs titer during subsequent biologic DMARDs therapy, including follow-up of anti-HBs titer and HBV DNA viral loads upon anti-HBs loss, to detect HBV reactivation earlier.

Ours is the first report that chronic kidney disease is a risk factor for loss of anti-HBs in patients receiving biologic DMARDs. This is an important issue because chronic kidney disease is prevalent among patients with rheumatic diseases nowadays, due to old age, diabetes-related nephropathy, and frequent use of nephrotoxic medications such as non-steroidal anti-inflammatory drugs or cyclosporine. Previous studies have shown that chronic kidney disease patients lose anti-HBs faster than do healthy subjects,[21, 22] and anti-HBs loss in chronic kidney disease or dialysis patients has been attributed to diminished interleukin-2 secretion, impaired macrophage function, decreasing memory B cell counts, and a weak amnestic response.[23–25]

This study had limitations. First, there are reports of increased likelihood of anti-HBs loss in patients with diabetes,[26, 27] and speculation that insulin resistance might affect T-cell differentiation and activation, and thereby cause immunologic dysfunction. Our relatively small sample size (four cases had diabetes mellitus) may explain why such an association was not evident. Second, despite considerable research into whether different biologic DMARDs equally increase the risk of anti-HBs loss or and HBV reactivation, results to date are inconclusive.[9, 28, 29] Only 15 cases were accrued and the small number precluded analysis of whether or not individual biologic DMARDs contributed equally to risk of losing anti-HBs.

### Conclusion

This hospital-based prospective study found that low baseline anti-HBs titer and chronic kidney disease independently predicted loss of anti-HBs in patients undergoing biologic DMARDs therapy to treat rheumatic diseases. This knowledge can be applied to identify patients at increased risk of becoming anti-HBs− and potential HBV reactivation from the onset of biologic DMARDs therapy. However, more research is needed to elucidate other risk factors for loss of anti-HBs and so refine the monitoring strategy to prevent HBV reactivation in patients receiving biologic DMARDs to treat rheumatic diseases.

## ACKNOWLEDGEMENTS

Dr David Neil, PhD, of Full Universe Integrated Marketing Ltd., Taiwan, provided professional editorial services. Ya-Chu Yang provided technical assistance in the laboratory work.

## Contributors

MHH made substantial contributions to the conception and design, analysis and interpretation of data, and drafting of the manuscript. YMC made substantial contributions to the conception and design, analysis and interpretation of data, and critical revision of the manuscript. YCT was involved in revision of the manuscript.

## Patient consent

Obtained.

## Ethics approval

Changhua Christian Hospital Institutional Review Board approved the study and all patients provided written informed consent for study participation.

## Provenance and peer review

Not commissioned; externally peer reviewed.

## REFERENCES

1. Perrillo RP, Gish R, Falck-Ytter YT. American Gastroenterological Association Institute technical review on prevention and treatment of hepatitis B virus reactivation during immunosuppressive drug therapy. Gastroenterology 2015;148:221–44.e3.

2. Torbenson M, Thomas DL. Occult hepatitis B. Lancet Infect Dis 2002;2:479–86.

3. Dervite I, Hober D, Morel P. Acute hepatitis B in a patient with antibodies to hepatitis B surface antigen who was receiving rituximab. N Engl J Med 2001;344:68–9.

4. Di Bisceglie AM, Lok AS, Martin P, et al. Recent US Food and Drug Administration warnings on hepatitis B reactivation with immune-suppressing and anticancer drugs: just the tip of the iceberg? Hepatology 2015;61: 703–11.

5. Shouval D, Shibolet O. Immunosuppression and HBV reactivation. Semin Liver Dis 2013;33:167–77.

6. Sarin SK, Kumar M, Lau GK, et al. Asian-Pacific clinical practice guidelines on the management of hepatitis B: a 2015 update. Hepatol Int 2016;10:1–98.

7. Tien YC, Yen HH, Li CF, et al. Changes in hepatitis B virus surface antibody titer and risk of hepatitis B reactivation in HBsAg-negative/HBcAb-positive patients undergoing biologic therapy for rheumatic diseases: a prospective cohort study. Arthritis Res Ther 2018;20:246.

8. Paul S, Dickstein A, Saxena A, et al. Role of surface antibody in hepatitis B reactivation in patients with resolved infection and hematologic malignancy: A meta-analysis. Hepatology 2017;66:379–88.

9. Charpin C, Guis S, Colson P, et al. Safety of TNF-blocking agents in rheumatic patients with serology suggesting past hepatitis B state: results from a cohort of 21 patients. Arthritis Res Ther 2009;11:R179.

10. Vassilopoulos D, Apostolopoulou A, Hadziyannis E, et al. Long-term safety of anti-TNF treatment in patients with rheumatic diseases and chronic or resolved hepatitis B virus infection. Ann Rheum Dis 2010;69:1352–5.

11. Hoofnagle JH. Reactivation of hepatitis B. Hepatology 2009;49(5 Suppl):S156–65.

12. Loomba R, Liang TJ. Hepatitis B Reactivation Associated With Immune Suppressive and Biological Modifier Therapies: Current Concepts, Management Strategies, and Future Directions. Gastroenterology 2017;152:1297–1309.

13. Chen YH, Chien RN, Huang YH, et al. Screening and management of hepatitis B infection in rheumatic patients scheduled for biologic therapy: consensus recommendations from the Taiwan Rheumatology Association. Formos J Rheumatol 2012;26:1–7.

14. Rothman KJ. Case-control studies. In: Rothman KJ, Greenland S, Lash TL, editors. Modern Epidemiology, Third Edition. Philadelphia: Williams & Wilkins; 2008. p. 111–27.

15. Suissa, S., Novel approaches to pharmacoepidemiology study design and statistical analysis. In: Strom B, editor. Pharmacoepidemiology, Fourth Edition. New York: John Wiley & Sons; 2005. p. 811–29.

16. Dixon WG, Kezouh A, Bernatsky, et al. The influence of systemic glucocorticoid therapy upon the risk of non-serious infection in older patients with rheumatoid arthritis: a nested case-control study. Ann Rheum Dis 2011;70:956–60.

17. Hung CH, Lu SN, Wang JH, et al. Correlation between ultrasonographic and pathologic diagnoses of hepatitis B and C virus-related cirrhosis. J Gastroenterol 2003;38:153–7.

18. European Association For The Study Of The Liver. EASL clinical practice guidelines: Management of chronic hepatitis B virus infection. J Hepatol 2012;57:167–85.

19. Koutsianas C, Thomas K, Vassilopoulos D. Hepatitis B Reactivation in Rheumatic Diseases: Screening and Prevention. Rheum Dis Clin North Am 2017;43:133–49.

20. Pei SN, Ma MC, Wang MC, et al. Analysis of hepatitis B surface antibody titers in B cell lymphoma patients after rituximab therapy. Ann Hematol 2012;91:1007–12.

21. Buti M, Viladomiu L, Jardi R, et al. Long-term immunogenicity and efficacy of hepatitis B vaccine in hemodialysis patients. Am J Nephrol 1992;12:144–7.

22. Tsouchnikas I, Dounousi E, Xanthopoulou, et al. Loss of hepatitis B immunity in hemodialysis patients acquired either naturally or after vaccination. Clin Nephrol 2007;68:228–34.

23. Sester U, Sester M, Hauk M, et al. T-cell activation follows Th1 rather than Th2 pattern in haemodialysis patients. Nephrol Dial Transplant 2000;5:1217–23.

24. Pesanti EL. Immunologic defects and vaccination in patients with chronic renal failure. Infect Dis Clin North Am 2001;15:813–32.

25. Descamps-Latscha B, Chatenoud L. T cells and B cells in chronic renal failure. Semin Nephrol 1996;16:183–91.

26. Leonardi S, Vitaliti G, Garozzo MT, et al. Hepatitis B vaccination failure in children with diabetes mellitus? The debate continues. Hum Vaccin Immunother 2012;8:448–52.

27. Joo EJ, Yeom JS, Kwon MJ, et al. Insulin resistance increases loss of antibody to hepatitis B surface antigen in nondiabetic healthy adults. J Viral Hepat 2016;23:889–96.

28. Tamori A, Koike T, Goto H, et al. Prospective study of reactivation of hepatitis B virus in patients with rheumatoid arthritis who received immunosuppressive therapy: evaluation of both HBsAg-positive and HBsAg-negative cohorts. J Gastroenterol 2011;46:556–64.

29. Papalopoulos I, Fanouriakis A, Kougas N, et al. Liver safety of non-tumour necrosis factor inhibitors in rheumatic patients with past hepatitis B virus infection: an observational, controlled, long-term study. Clin Exp Rheumatol 2018;36:102–9.

